# Rewilding alters mouse epigenetic aging

**DOI:** 10.1101/2024.09.25.615023

**Authors:** Matthew N Zipple, Ivan Zhao, Daniel Chang Kuo, Sol Moe Lee, Michael J Sheehan, Wanding Zhou

**Author notes:** Authors contributed equally (Matthew N Zipple, Ivan Zhao). **AVAILABILITY:** The generated mouse methylome profiles (N=93) are available in the Gene Expression Omnibus with accession GSE269932. Other public lab mouse liver methylomes (N=7) and additional samples for epigenetic clock construction can be found in GSE184410 (sample accessions listed in Table S1). Informatics for mouse methylation data preprocessing and functional analysis is available in the R/Bioconductor package *SeSAMe* (version 3.22+): https://bioconductor.org/packages/release/bioc/html/sesame.html. **DECLARATION OF INTERESTS** The authors declare no competing interests.

## Abstract

The aging of mammalian epigenomes fundamentally alters cellular functions, and such changes are the focus of many healthspan and lifespan studies. However, studies of this process typically use mouse models living under standardized laboratory conditions and neglect the impact of variation in social, physical, microbial, and other aspects of the living environment on age-related changes. We examined differences in age-associated methylation changes between traditionally lab-reared and “rewilded” C57BL6/J mice, which lived in an outdoor field environment with enhanced ecological realism. Systematic analysis of age-associated methylation dynamics in the liver indicates a genomic region-conditioned, faster epigenetic aging rate in mice living in the field than those living in the lab, implicating perturbed 3D genome conformation and liver function. Altered epigenetic aging rates were more pronounced in sites that gain methylation with age, including sites enriched for transcription factor binding related to DNA repair. These observations underscore the overlooked role of the social and physical environment in epigenetic aging with implications for both basic and applied aging research.

## MAIN TEXT

Much biomedical research is conducted on model organisms living under standardized laboratory conditions^1^. Unlike natural populations, model organisms living in lab settings have stable access to food and shelter, experience mild, near-constant climatic conditions, and often have limited, static social experiences^2^. One consequence of keeping animals in the lab is that median lifespans are dramatically extended, partially through reduced extrinsic mortality from predators and competition (similar to lifespan comparisons in zoo versus wild populations^3^). However, laboratory conditions may also influence intrinsic cellular processes such that the magnitude of age-related molecular changes differ between laboratory and field conditions, with downstream consequences for senescence.

Changes in DNA methylation are a molecular hallmark of aging^4,5^. Global methylation tends to decline with age while specific CpGs, *e.g*., at CpG islands, become more methylated with age^6^. This age-associated methylation change is the basis of epigenetic clocks—inference models that predict age from the methylation profile^7^. The aging epigenome also mechanistically contributes to other cellular and physiological changes linked to aging^4^, such as stem cell functional deficit, malignancy risk, etc.

The extent to which age-associated epigenetic change is shaped by environmental context and whether the changes observed under laboratory context accurately generalize to more ecologically realistic conditions remain open questions. Recently, a study of house mice living in the wild identified increased age-associated rates of change in DNA methylation patterns in a handful of loci compared to laboratory controls^8^, suggesting either that wild genotypes or free-living conditions may increase rates of epigenetic change in mice. Direct comparisons of epigenetic aging of the same genotype under lab versus field conditions are needed to identify the role of lab environments in shaping epigenetic aging patterns.

We employed a “rewilded” model of mouse research to address this question. We released the common lab mouse (strain C57BL6/J) into a controlled outdoor field environment (Figure 1A, S1A), which offers dramatically increased social and physical ecological realism compared to standardized lab conditions^2^. This approach allows us to retain lab models’ manipulability and reproducibility while allowing for dynamic and complex physical and social experiences. We collected 73 liver tissue samples from rewilded mice 26-302 days of age (Figure 1B) that had lived in the field enclosure for a variable number of days (Figure S1A-B). We then compared the liver methylomes of these rewilded samples with those collected from 27 C57BL6/J mice of similar age ranges that lived their whole lives in a standard lab colony (Figure 1B).

**Figure 1:**
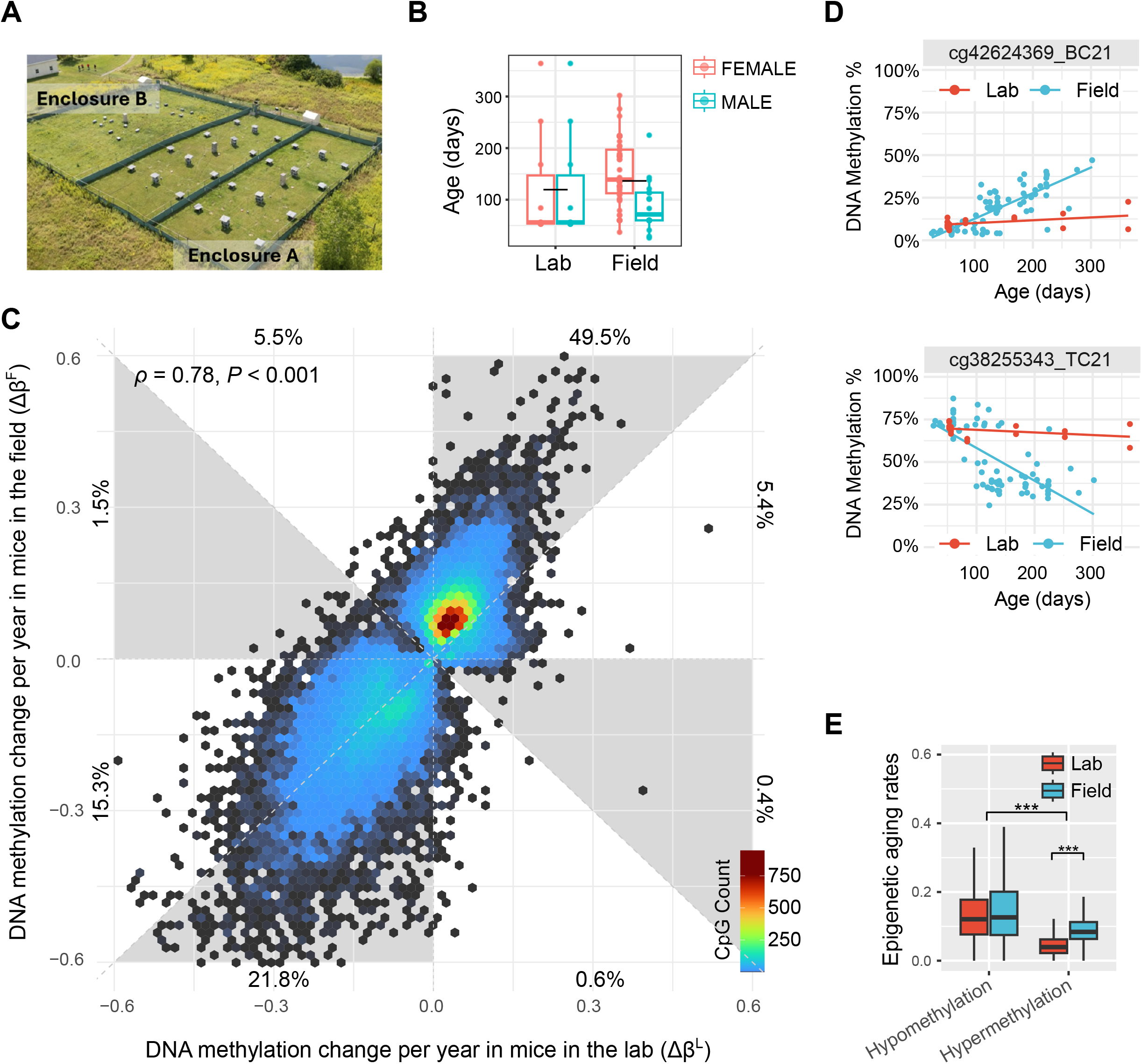
Comparison of age-related epigenetic changes in lab vs field environments. (A) An aerial view of the two enclosures where the field cohort lived (B) Field and lab cohorts were broken down by environment, age, and sex (color). (C) Comparing epigenetic aging rates between lab (X-axis) and field mice (Y-axis) among those sites that show age-associated hyper- or hypo-methylation in mouse liver tissues. The rates represent the estimated change in methylation level per year. The percentages of sites falling into the eight evenly spaced angular octants are labeled next to the axes. (D) Representative CpGs with faster epigenetic aging rates in the field than in the lab (upper panel: age-associated hypermethylation; lower panel: age-associated hypomethylation). (E) Boxplots compare the distributions of absolute annual rates of epigenetic change at sites that become hypo- and hyper-methylated in each environment.

To measure the environmental impact on rates of change in methylation, we modeled cytosine methylation levels (percentage cytosine modified) at ∼277K CpG sites (from ∼285K after filtering missing values) using age, sex, and living environment (laboratory or field) as predictors. We explicitly modeled the interaction between age and living environment to evaluate environment-specific epigenetic aging rates (the coefficient of the age slope). Analysis of CpGs showing age-associated methylation change in either cohort (q<0.01) suggests lab and field mice are moderately congruent in the annual rate of methylation change (Spearman’s rho = 0.78, *P*-value < 0.001, N = 33,038), indicating site-specific intrinsic propensities of the DNA methylation drift (Figure 1C). Representative CpGs with similar rates of methylation aging, faster aging in the field than in the lab, and vice versa, are shown in Figure 1D and Figure S1C-D.

Depending on the estimated rates of methylation change in animals living in the lab and the field, we categorized age-associated methylation changes into eight octants in Figure 1C. 54.9% and 37.1% of the age-associated methylations are bi-cohort joint hypermethylations and hypomethylations, respectively. In contrast, 7% of CpGs show hypermethylation in the field but hypomethylation in the lab. Only ∼1% of CpGs show hypomethylation in the field but hypermethylation in the lab. Age-associated hypomethylations tend to have larger effect sizes than hypermethylations (Figure 1E), consistent with a global decline in methylation levels with age.

Age-associated increases in methylation are notably faster in mice living in the field than in the lab (Figure 1C). Of age-associated joint hypermethylations, 90.2% (49.5% in 54.9%) have faster epigenetic aging rate estimates in the field. Among sites that show significant (q < 0.01) increases in methylation in both environments, the average rate of epigenetic change is approximately 45% faster in the field (95% CI = 42%-48%). In contrast, only moderate acceleration is found in age-related hypomethylations. 59% (21.8% in 37.1%) of the joint hypomethylations have faster aging rates in the field. Restricting our analysis to sites that display a significant (q < 0.05) age-by-environment interaction yields the same conclusion—many more sites display faster hypermethylation in the field than in the lab (545/555, 98.2%) while comparable numbers of sites display faster hypomethylation in the field versus the lab (43% are faster in the field, 57% faster in the lab). This intriguing disparity between hyper- and hypo-methylation suggests that different mechanisms of aging may be variably affected by environmental differences.

To explore this difference, we performed a functional enrichment analysis. Broadly, sites that displayed congruent patterns of hyper- and hypomethylation under both environments reflect previously reported patterns of age-associated methylation change. Those sites that display substantial hypermethylation with age in both environments primarily localize to bivalent promoters, poised enhancers, and binding sites of the Polycomb repressive complexes (*e.g*., SUZ12, MTF2, AEBP2, JARID2, CBX7, PCGF2, see Figure 2A), consistent with prior reports of their association with replicative epimutation^9^ and role in stem cell differentiation^10,11^. Sites that became substantially hypomethylated with age in both environments localize to the binding sites of transcription factors key to hepatocyte development and function (*e.g*., PROX1, NCOR1, ONECUT1, NR5A2, NFIB, see Figure 2B), suggesting the role of these hypomethylation events in age-associated liver function maturation. Joint hypomethylations are also linked to the binding of cohesin complex proteins (*e.g*., SMC3, SMC1A, RAD21, STAG2) and CTCF. These scaffold proteins involved in 3D nuclear conformation (Figure 2B) reflect previously reported age-associated chromatin architectural changes^12^. As an additional confirmatory control, sex-associated methylations are exclusively found at sex chromosomes and DMC1 binding (Figure S2A).

**Figure 2:**
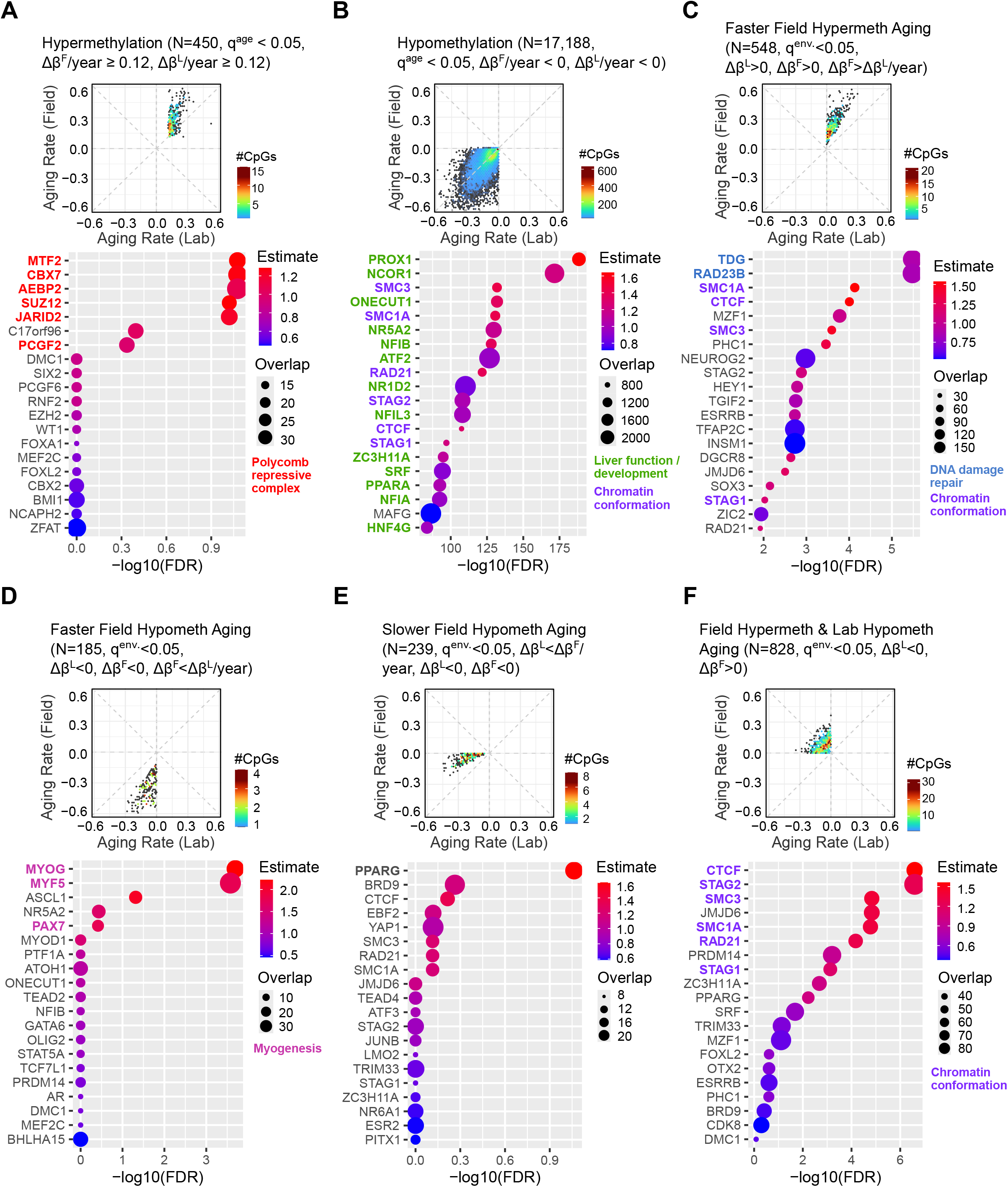
Functional enrichment analysis of age-associated methylations with congruent and discrepant rates between mice living in the lab and the field. Transcription factor binding sites enrich (A) methylations that gain with age in both lab and field-living mice; (B) methylations that lose with age in both lab and field-living mice; (C) methylations that gain faster with age in the field than lab mice; (D) methylations that lose faster in the field than the lab-living mice; (E) methylations that lose faster in the lab than field mice; (F) methylations that gain in the field but lose with age in the lab-living mice. Transcription factor binding peaks were obtained from mouse ENCODE databases and overlapped with the mouse methylation array probes. CpG sites with age-associated q-values smaller than 0.05 were used as queries.

Sites that gain methylation faster with age in the field are enriched for binding by two DNA damage repair genes, TDG and RAD23B (Figure 2C). This enrichment suggests that the faster hypermethylation observed in the field mice is linked to age-related DNA damage^13^, epigenetic silencing^14^, and repair. Increased DNA damage and its repair is a molecular hallmark of aging^5,15^, and our results indicate that the rate of epigenetic change linked to this process depends on environmental conditions. Sites that gain methylation more rapidly in the field are also enriched for DNA binding of several genes involved in chromatin conformation (SMC1A, CTCF, SMC3, STAG1), likely reflecting an age-associated 3D genome conformation degradation. Few sites gained methylation more rapidly in the lab than in the field, without any significant enrichment of binding sites associated with these changes.

Sites that lose methylation more rapidly in the field are moderately associated with transcription factors conventionally involved in the development of non-hepatocytes, such as muscle cells (MYOG, MYF5, PAX7) and neurons (ASCL1) (Figure 2D, S2B). Whether this is due to tissue composition change or altered hepatocyte expression warrants further investigation. Assessing differences in rates of methylation change in other tissue types represents a rich opportunity to understand tissue and system-specific environmental impacts on the aging process. Sites that lost methylation more rapidly under laboratory conditions did not show a strong signal in the enrichment analysis (Figure 2E), with the only significant binding enrichment observed being PPARG. This gene regulates lipid storage in the liver^16^.

Next, we examined sites exhibiting divergent directions of change in epigenetic aging patterns between the lab and the field. While both cohorts exhibited common methylation loss at cohesin complex and CTCF binding sites (Figure 2B), mice in the field uniquely gained additional methylation at a distinct subset of CpGs bound by these same complexes (Figure 2F). This observation aligns with the understanding that CTCF binding is highly responsive to DNA methylation, contingent on genomic and chromatin context^17^. Our findings suggest that 3D chromatin conformation changes likely drive epigenetic aging and contribute to the differential rates of epigenetic change observed between the lab and the field. Unlike sites that display hypomethylation with age in both environments, no strong involvement of liver function and development is observed in these sites with divergent directions of age-associated changes in methylation (Figure 2F).

Finally, we asked if the observed disparity in rates of epigenetic change is reflected in epigenetic clock measurements. Epigenetic clocks are predictive models for chronological and phenotypic age based on methylation levels or other molecular markers^18,19^. We employed an epigenetic clock of 248 CpG features to predict the age of each mouse tissue sample (Figure S2C, Methods). The clock overpredicts age more in the field than in the lab. The median residual of the predicted ages for the field (2.14 months) was slightly higher than that of the lab (1.44 months), respectively, with a marginal *P*-value of 0.0764 (Figure S2D). Despite clocks using a small fraction of CpG features in the genome, the higher prediction residuals confirm the above observation of differential epigenetic aging in the field versus the lab. Yet, relying only on the single quantitative output of such a clock conceals the variation in the magnitude of the environmental effects described above, which depend on whether sites become hyper versus hypo methylated with age. When designing clocks in the future, our results suggest that researchers who aim to detect environmental impacts on rates of epigenetic change should bias their selection process towards sites that gain methylation with age rather than lose it.

Our results lead us to two conclusions. First, the rate of epigenetic changes in the most used biomedical model organism is highly dependent on environmental context. Second, this more rapid aging of the epigenome is particularly pronounced in sites that gain methylation with age, which are enriched for genes associated with DNA damage repair and CTCF and cohesin binding. From the current data, it remains unclear if rewilded mice would also show accelerated senescent phenotypes, including early onset disease development and behavioral declines, compared to those in the lab.

Mice hold a central position in aging research. Our results highlight the need to consider the environmental context of model organisms when studying aging-related phenotypes. Although therapeutics targeted at physiological and behavioral decline are usually tested on mice living in standard conditions, these mice are physiologically distinct from the same genotype, experiencing more ecologically realistic conditions. Both mice and humans face fluctuating physical and social environments in natural populations. Our results demonstrate the importance of this variation in the molecular mechanisms of aging.

## ACKNOWLEDGMENTS

This work is supported by Pilot and Feasibility awards to MNZ and MJS from the Animal Models for the Social Dimensions of Health and Aging Network (project #5R24AG065172-03) and the National Institute of Health [R35-GM146978] to WZ. MNZ was also supported by a Klarman Postdoctoral Fellowship award from Cornell University and an NSF postdoctoral fellowship in biology (award # 2109636). We thank Brian H. Chen for providing mouse methylation arrays, and Maria Lemma from the CHOP CAG core for the array experiments. We also thank Jenny Tung for valuable conversations and feedback on this project.

## AUTHOR CONTRIBUTIONS

This project was first conceptualized by MNZ and MJS, and field mouse liver samples were provided by MNZ, DCK, and MJS and profiled by SML. All participated in the data analysis. MNZ, MJS, and WZ oversaw the project.

## METHODS

### Field enclosures and study subjects

The enclosures at Cornell University’s Liddell Field Station were described in detail previously^20^. Here, we only describe elements critical to this experiment. Across the trial, we allowed animals to live freely in two outdoor field enclosures for portions of their lives (Figure S1A-B).

We generated our study subjects by breeding nine-week-old male and female C57BL/6J mice that we purchased from Jackson Laboratory (Bar Harbor, ME) and mated in our laboratory at Cornell University. Upon female pregnancy, the males were removed from breeding cages to prevent re-insemination following parturition. When pups were 8-10 days of age, we anesthetized litters and their mothers using isoflurane and injected either 1 (pups) or 2 (mothers) RFID tags (Biomark Mini HPT10) subcutaneously. The RFID tag is a permanent identification method to identify known-aged individuals following re-capture from our field enclosures. Our field study subjects (n = 73 total) were a combination of these mothers and pups. Individuals spent varying amounts of time living in the laboratory and our field enclosures (Figure S1A). A portion of the mice examined here contributed to other behavioral datasets during their ontogeny^21^.

We released most of our study subjects (Group 1; n = 41) and their littermates and mothers into our field enclosures at 12-16 days of age. These mothers comprise an additional subset of our samples (Group 2; n = 11). Until Group 1 animals were ∼60 days of age (or the end of their life, whichever came first), they lived in an enclosure (Enclosure A) 15m x 38m in size, approximately 9,000 times the area of a typical mouse cage. In this enclosure, we set up 16 plastic tubs (31-gallon storage totes, Rubbermaid, USA), placed into four neighborhoods of four resource zones (Figure S1B). Each tub (hereafter “resource zones”) contained ad libitum food access and a nestbox that provided insulation and shelter from adverse weather conditions. We equipped each zone with a single joint entrance/exit made from a 6-inch-long PVC pipe (2 inches in diameter). These resources and the single entrance made the resource zones highly valuable and defendable, mimicking commensal mice’s foraging landscape. Group 2 mothers lived alongside their Group 1 pups as they developed.

When pups were ∼60 days of age, all Group 1 and Group 2 animals were re-captured. A subset of these animals were then sacrificed (Group 1, n = 10; Group 2, n = 6). The remaining animals were transferred to another enclosure (Enclosure B). This transfer allowed us to continue to use Enclosure A, which contains more intensive RFID and camera tracking technology than Enclosure B, for other experiments. The remaining Group 1 and Group 2 animals continued to live out their lives in Enclosure B. Enclosure B is approximately 1.5 times the size of Enclosure A, and we set up resource zones in Enclosure B in the same pattern as Enclosure A.

At this time, we also introduced an additional cohort of animals into Enclosure B. This cohort of animals was made up of (1) individuals of the same age as Group 1 but who had spent their lives under standard laboratory conditions (hereafter Group 3; n = 12) and (2) the mothers of the individuals in Group 3 (hereafter Group 4, n =10), who were purchased and mated at the same time as the mothers in Group 2. Thus, there was some variation in the total amount of time and the extent of individuals’ lives that animals in the ‘field environment’ spent living in our enclosures. However, all animals spent a substantial amount of time living in the field environment (median proportion of life spent in the field = 0.79, min = 0.16, max = 0.94; Figure S1B).

### Rewilded mice liver methylome profiling

Tissues came from one of two sources. A small number (n = 4) were collected from animals that we re-collected from the field enclosure before age 60 days and who had lost their RFID tags. These animals were of known age and sex but unknown identity and were sacrificed. A larger number (n = 11 Group 1, n= 5 Group 2) were captured via live Sherman traps placed overnight in Enclosure A when Group 1 animals were approximately 8.5 weeks old (and Group 2 animals were approximately 20 weeks old). Most samples (n = 58) came from animals from Groups 1-4 while they were living in Enclosure B. On an approximately fortnightly basis, we placed Sherman live traps in Enclosure B and sacrificed animals we caught overnight. During all sample collection, animals were humanely euthanized via cervical dislocation followed decapitation. We then collected one liver lobe and flash-froze the tissue on dry ice. Tissues were then stored at -80 C until DNA extraction. We extracted DNA from 25 mg of liver tissue, following the DNeasy Blood and Tissue Kit for DNA Extraction protocol (Qiagen N.V.). We stored the extracted DNA at -80C until methylome profiling.

According to the manufacturer’s protocol, bisulfite conversion of 500 ng input liver DNA per sample was performed using EpiTect Bisulfite Kits (Qiagen, 59104). The Infinium Mouse Methylation BeadChip assays were conducted at the Center for Applied Genomics Genotyping Core of the Children’s Hospital of Philadelphia.

### Lab mouse liver methylome profiling

C57BL/6J mice were acquired from the Jackson Laboratory. Extraction of genomic DNA from mouse liver tissues follow previous work^22^. The whole liver tissue was homogenized using a tissue homogenizer (OMNI, TH115) in 500 μl of lysis buffer containing 10 mM Tris pH 8.0 (VWR, 97062-674), 300 mM NaCl (VWR, 10128-484), 0.5% SDS (Invitrogen, 15553027), and 5 mM EDTA (VWR, 10128-442). After adding 15 μl of Proteinase K (NEB, P8107S), the solution was incubated at 55°C overnight. Subsequently, 100 μl of the solution was combined with an additional 400 μl of lysis buffer and incubated at 55°C for 2 hours. The 500 μl of the solution was placed into a 5PRIME Phase Lock Gel tube (Quanta bio, 10847-802) pre-centrifuged for 1 minute. 500 μl of phenol/chloroform/isoamyl alcohol (Sigma-Aldrich 77617) was added to the phase Lock Gel tube. The aqueous phase solution was transferred to a new 1.5 mL centrifuge tube (Eppendorf, 05414203). 500 μl of 100% isopropanol (MilliporeSigma, EM1.09634.1011), GlycoBlue (Invitrogen™, AM9515), and Ammonium acetate solution 7.5M (Sigma-Aldrich, A2706) were added to the tube. The solution was vortexed and incubated at -20°C for 30 min up to O/N. Samples were centrifuged at 16,000g for 30 min at 4°C and washed twice by adding 1 ml 70% EtOH (MilliporeSigma, EM1.00983.1011). After the last wash and removal of 70% EtOH, the samples were air-dried for 10 min and resuspended in 200 μl of the Tris buffer pH8.0 (VWR, 97062-674), and incubated at 55°C for 10 min. DNAs that were not completely dissolved were incubated at 4°C for overnight. If the dissolved DNA did not exhibit a transparent color or the DNA quantity was inadequate, an additional bead purification step was carried out. Briefly, 2x volume of AMPure XP (Beckman Coulter, A63881) was added to the DNAs, mixed thoroughly, briefly spun down, incubated the mixture for 5 min at room temperature, placed on a magnet stand, and washed 2 times with 500 μl of freshly made 80% EtOH, allowed to dry for 3 to 5 mins. The final elution was performed using 100 ∼ 200 μl autoclaved ultrapure water. DNA amount was measured using the Qubit 4 Fluorometer (Invitrogen) with the dsDNA HS Assay Kit (Invitrogen, Q33231).

DNA bisulfite conversion was performed using the EZ DNA Methylation Kit (Zymo Research, D5001) or EZ-96 DNA MethylationTM MagPrep (Zymo, D5040). Samples with bisulfite converted by EZ DNA Methylation kit were performed according to the manufacturer’s instructions with the specified Illumina Infinium Methylation Assay modifications. Samples that bisulfite converted by EZ-96 DNA MethylationTM MagPrep (Zymo, D5040) were performed using the same process as the above. The Infinium Mouse Methylation BeadChip assays were conducted at the Center for Applied Genomics Genotyping Core of the Children’s Hospital of Philadelphia.

### Public lab mouse liver methylome dataset

We also included 10 C57 lab mouse liver samples from a prior study (GSE184410; see Supplemental Table S1 for sample accessions). IDAT files were downloaded and processed using the openSesame workflow with default parameters.

### Data preprocessing and analysis

All IDAT files were processed using the openSesame workflow with default parameters^23^. The data quality is assessed using the sesameQC pipelines. All samples have over 90% probes passing the signal detection threshold (pOOBAH detection *P*-value < 0.05).

### Construction of the epigenetic clock

We collected 706 public mouse tissue methylomes (Supplemental Table S1) and constructed an epigenetic clock for the MM285 array using an elastic net framework. The elastic-net regularized linear model was built using glmnet. To select the most predictive CpGs, we set alpha to 0.5 and lambda to 0.098, selected using the cv.glmnet function, which automatically optimizes the mean absolute error of the model using 10-fold cross-validation. This procedure leads to an epigenetic clock of 248 CpG probes with an estimated mean absolute error of 1.2 months.

---

**Supplemental Figure S1:**
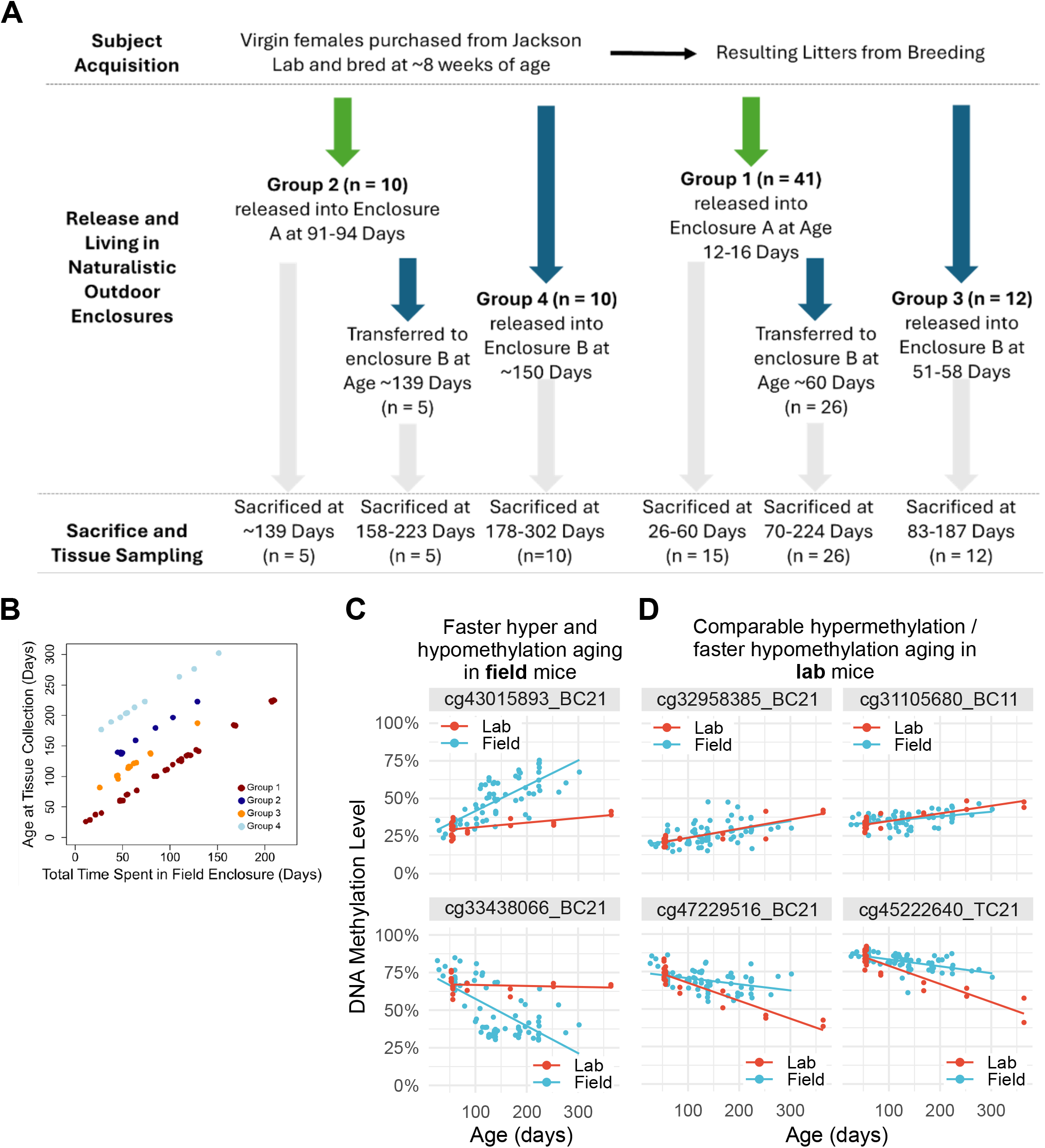
(A) Schematic of the life experiences of the subjects in this study. All animals (N=73) spent some time under laboratory and semi-natural conditions. Most individuals (Group 1, n = 41) spent nearly all their lives outside under semi-natural conditions. (B) Scatter plot of individuals’ ages at the time of tissue collection (y-axis) versus the time spent in the enclosures (x-axis). Points are jittered to prevent overlap. N=73 tissue samples are plotted. (C-D) Representative CpGs that display differential epigenetic aging, comparing age in days (X-axis) and methylation level (Y-axis) in faster aging in field mice (C) and lab mice (D). Top row shows comparable or faster hypermethylation and bottom row shows faster hypomethylation.

**Supplemental Figure S2:**
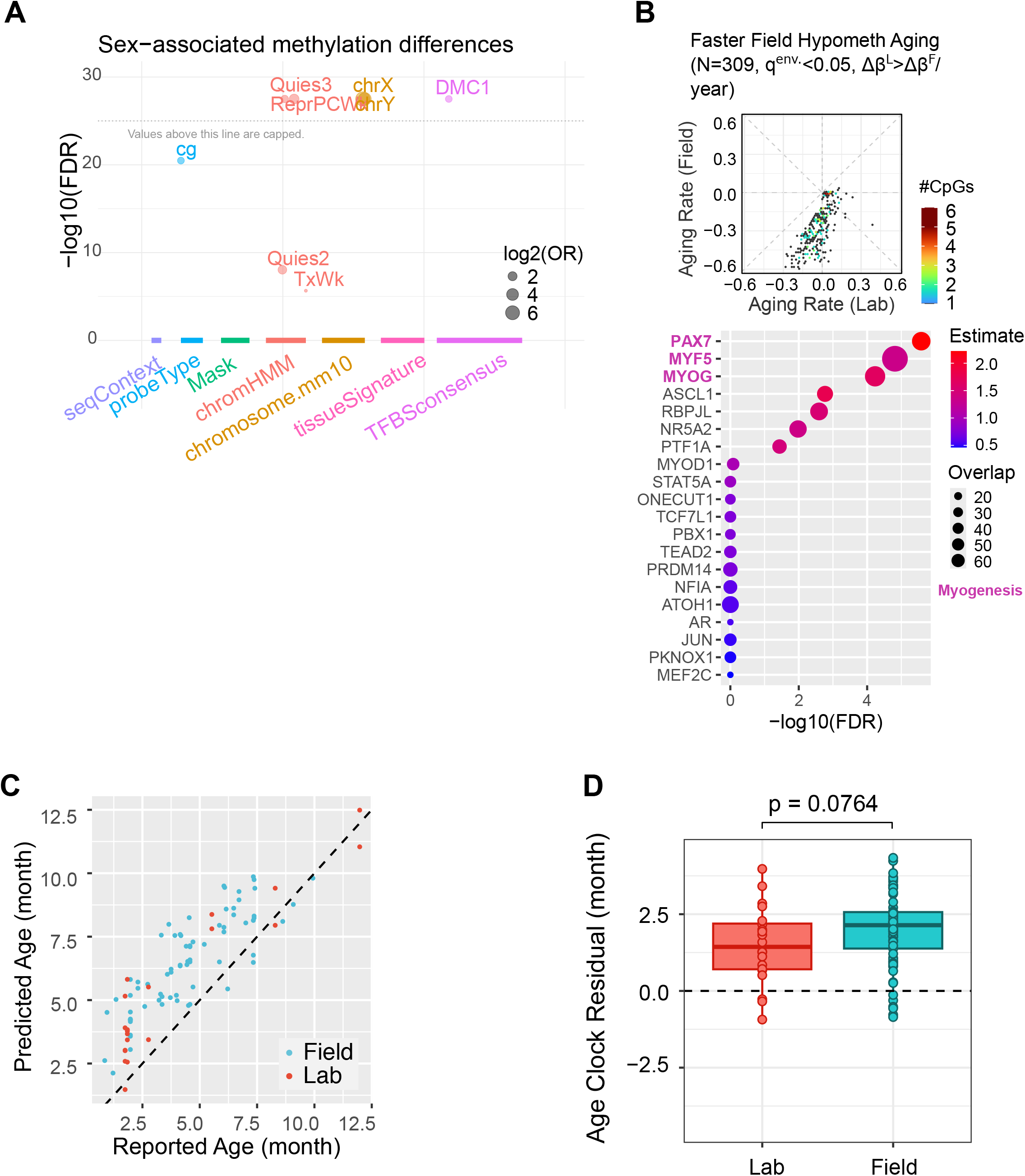
(A) Enrichment of sex-associated methylation differences. (B) Functional enrichment analysis of age-associated methylations that lose faster in the field than the lab-living mice; This is the same as Figure 2D except that no directionality is restricted, hence including a small number of sites that gain methylations in the lab-living mice. (C-D) Epigenetic clock reveals accelerated biological aging in field mice using 248 CpG methylation features. (C) Scatter plot comparing clock predictions in lab and field mice. (D) Box plot comparing epigenetic clock prediction residuals in lab and field mouse cohorts. Wilcoxon rank sum test, *p* = 0.0764.

